# Molecular and genetic heterogeneity in iPSCs derived from an outbred laboratory mouse population

**DOI:** 10.64898/2026.05.02.722403

**Authors:** Madison Armstrong, Anne Czechanski, Emily Swanzey, Qiongyu Chen, Whitney Martin, Callan O’Connor, Catherine Brunton, Selcan Aydin, Hannah B. Dewey, Steven C. Munger, Laura G. Reinholdt

## Abstract

Genetically diverse panels of human pluripotent stem cells enable genetic dissection of cellular phenotypes, but comparable induced pluripotent stem cell (iPSC) resources in model organisms remain limited. We generated a panel of iPSCs from the Diversity Outbred (DO) mouse population and established 288 genetically unique lines that retain the allele frequency distribution, heterozygosity, and low population structure of the source population. The lines exhibit consistent growth, pluripotent gene expression profiles, and capacity to form embryoid bodies. Transcriptomic profiling of the lines revealed significant variation in gene expression driven in part by genetic background. We used expression quantitative trait locus (eQTL) mapping to identify over 10,000 regulatory loci that influence gene expression variation, including multiple distal eQTL hotspots that are key gene regulatory hubs and are shared with DO embryonic stem cells (ESCs). The largest hotspot, mediated by *Lifr*, showed a shift in founder allele effects relative to ESCs, consistent with differences in cellular state and culture conditions. These results establish the DO iPSC panel as a genetically diverse, publicly accessible platform derived from a laboratory mouse genetic reference population, enabling integration of *in vitro* cellular phenotypes with *in vivo* traits within a closed, outbred population.

## INTRODUCTION

Pluripotent stem cells (PSCs) including embryonic stem cells (ESCs) and induced pluripotent stem cells (iPSCs) derived from model organisms have well-established utility for genetic engineering, and more recently, for *in vitro* systems genetic analysis, which utilizes panels of cell lines derived from genetic reference panels. Mouse genetic reference panels consist of inbred strains, as well as derivative recombinant inbred and outbred populations that are used for iterative phenotyping, genetic mapping, and experimental manipulation [1–3]. For example, the Collaborative Cross (CC) panel was bred from 5 phenotypically distinct common laboratory inbred strains (A/J, C57BL/6J, 129S1/SvImJ, NOD/ShiLtJ, NZO/HlLtJ) and 3 wild-derived inbred strains (CAST/EiJ, PWK/PhJ, and WSB/EiJ). The resulting CC panel is a collection of recombinant inbred (RI) strains that capture haplotypes from these 8 “founder” inbred strains and enable high genetic reproducibility [4]. The Diversity Outbred (DO) mouse population was created through random outcrossing of partially inbred CC RI lines and features high heterozygosity, balanced founder allele frequencies, and recombinant chromosomes that facilitate high resolution genetic mapping [5, 6].

The DO population exhibits high phenotypic diversity across a range of heritable, quantifiable traits that govern behavior [7], physiology [8–10], disease susceptibility [11, 12], environmental / drug response [13, 14], and others. This phenotypic diversity is also observed *ex vivo,* for example in pancreatic beta cells (insulin secretion) [11], mouse ESCs (mESCs) (LIF response, differentiation capacity) [15–18], and in fibroblasts (morphology, heavy metal response traits) [19]. By harnessing the genetic architecture of the DO, in combination with epigenomic, transcriptomic, and proteomic data [15, 18, 20, 21], the genetic regulation of these molecular and cellular traits can be exposed and further anchored to disease-relevant physiological traits in genetically-matched mice [22, 23]. In laboratory mice, this iterative *in vivo / ex vivo* genetic analysis and experimental validation can be enabled by publicly available resources of mouse strains with matched panels of cells [24].

Human iPSCs derived from diverse populations are now widely used for genetic analysis of cellular phenotypes, and comparable resources from model organisms are emerging. In addition to mESCs from a variety of inbred strains [25] and recombinant inbred strains (e.g. BXD, [15]), there is one publicly available resource of genetically heterogeneous DO mESCs [18] and one resource of DO epiblast stem cells (EpiSCs) [16], as well as induced pluripotent stem cells (iPSCs) from a variety of inbred strains [26, 27], but there is no publicly available panel of iPSCs from the Diversity Outbred population. As with many cellular traits, strain genetic background has been shown to influence the derivation efficiency, stability, and differentiation capacity of pluripotent stem cells whether they are derived from mice [15–18, 21, 22, 26–28] or humans [29–31], so they are attractive models for studying the genetic regulation of early development and cell fate.

To address this gap, we created and characterized a panel of iPSCs from the DO mouse population. The panel preserves the allele frequency distribution and heterozygosity of the source population and supports high-resolution genetic mapping. Using this resource, we performed expression quantitative trait locus mapping and identified regulatory loci that influence transcriptional variation across lines. Comparison with DO embryonic stem cells revealed conserved regulatory hotspots alongside shifts in founder allele effects consistent with differences in culture conditions.

## RESULTS AND DISCUSSION

We randomly selected 329 tail tip fibroblast lines that were previously derived from several cohorts of DO mice, favoring lines with low passage stocks (generations 34-37, IMSR_JAX:009376) [19] and balanced representation of male and female lines. We reprogrammed these lines through lentiviral delivery of an inducible, polycistronic cassette containing the four murine reprogramming genes, *Oct4*, *Sox2*, *Klf4*, and *cMyc* (OKSM). During this process we found that a subset of the fibroblast lines (n = 33) grew poorly or failed to reprogram. The remaining 296 polyclonal iPSC lines were frozen at P3, genotyped, and expanded to P5-P6 for QC and biobanking at The Jackson Laboratory (Fig 1, Supplemental Table 1). We flagged 8 lines from the core panel because they shared a common animal ID or were later found to have unexpectedly high kinship (see *Genetic architecture* below and Supplemental Table 1), yielding a total of 288 genetically diverse, non-redundant DO iPSC lines. We also flagged 4 lines where the genotyped sex (XX, XY) did not match the sex of the animal from which the source tissue was derived (Supplemental Table 1). Supplemental Table 1 delineates the iPSC lines with associated metadata (including source fibroblast line IDs and source animal IDs) and bankstock information that can be used to obtain lines from The Jackson Laboratory.

**Figure 1.**
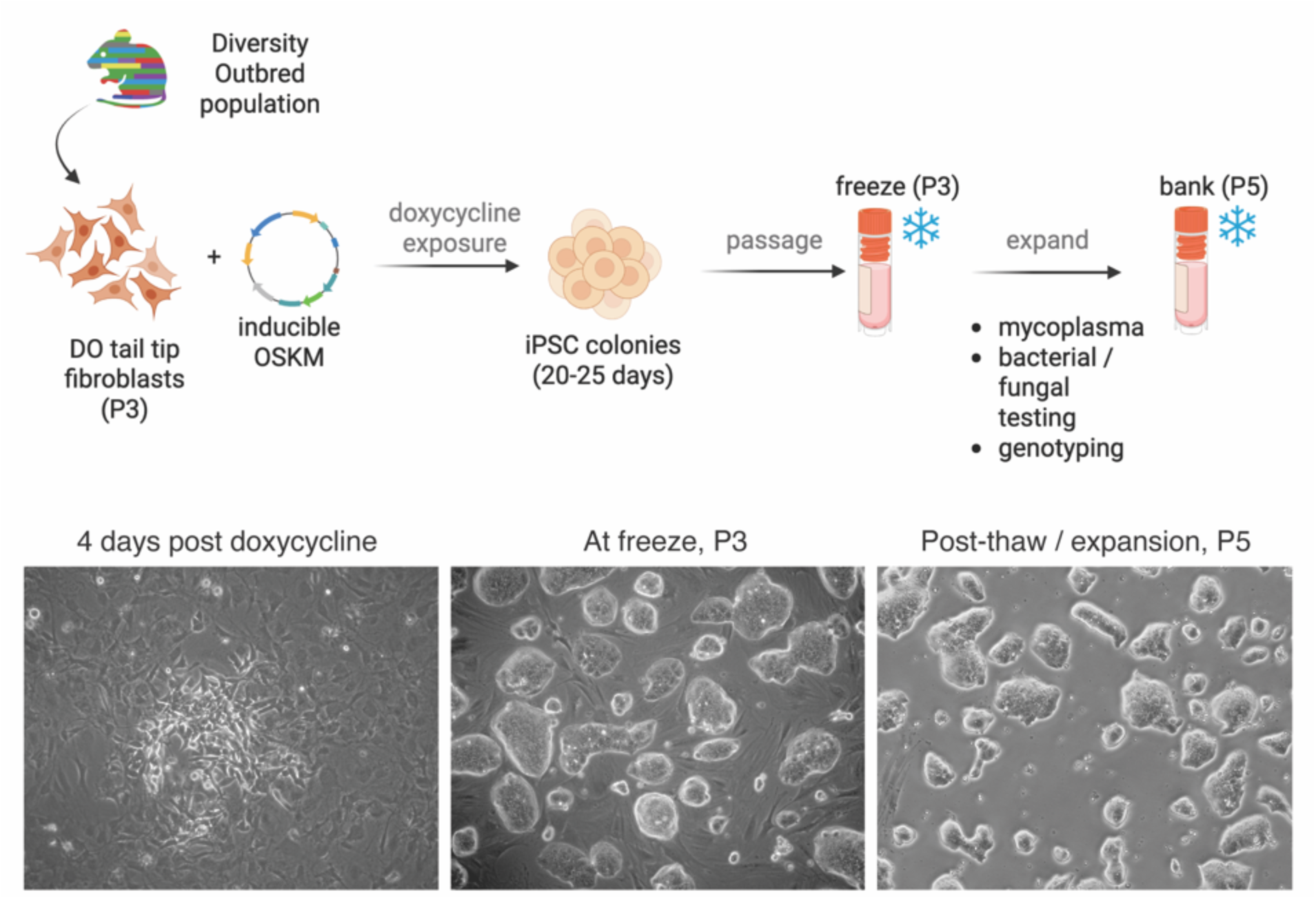
Overview of DO fibroblast reprogramming and iPSC derivation. Tail tip fibroblasts from non-sibling Diversity Outbred mice were cultured on gelatin-coated plates and reprogrammed by lentiviral transduction of a doxycycline-inducible mouse OKSM cassette. Twenty-four hours after transduction, virus-containing medium was replaced with mESM + 2i/LIF supplemented with doxycycline to induce expression of the reprogramming factors. Emerging iPSC colonies were pooled and expanded as polyclonal lines. After three passages, lines were cryopreserved at P3. One vial per line was subsequently expanded to P5 for quality control screening and genotyping. Finalized lines were banked with associated metadata in The Jackson Laboratory Biobank (see Supplemental Table 1).

### Genetic architecture

The GigaMUGA genotyping array consists of probes for 141,090 single nucleotide polymorphisms (SNPs) genome-wide that were specifically chosen to discriminate among the 8 founder inbred strains of the DO/CC populations [32]. SNP-level genotypes from the GigaMUGA were used to infer 8-state founder allele probabilities, which in turn were used to reconstruct the 36-state diplotypes of each DO iPSC line. An important feature of the DO population is that, at any locus, the relative frequency of alleles from each of the 8 founder inbred strains are generally balanced across the population (i.e., Minor Allele Frequency [MAF] ≥ ⅛ or 0.125) due to the large breeding population and random outcrossing scheme, which provides higher genetic mapping power with modest sample sizes compared to natural breeding populations [33]. We calculated genome-wide allele frequency and found a modest genome-wide deviation from the 1/8 expectation (χ2 (7) = 22.64, p = 0.0020) — driven mostly by under-representation of PWK/PhJ and over-representation of NOD/ShiLtJ alleles — though absolute shifts overall were small (∼1-1.5%; Fig 2A, Supplemental Table 2). We could not attribute loss or gain of any founder strain haplotype to reprogramming efficiency from these results, though genetic background has been previously correlated with reprogramming efficiency of inbred strain mouse embryonic fibroblasts [27].

**Figure 2.**
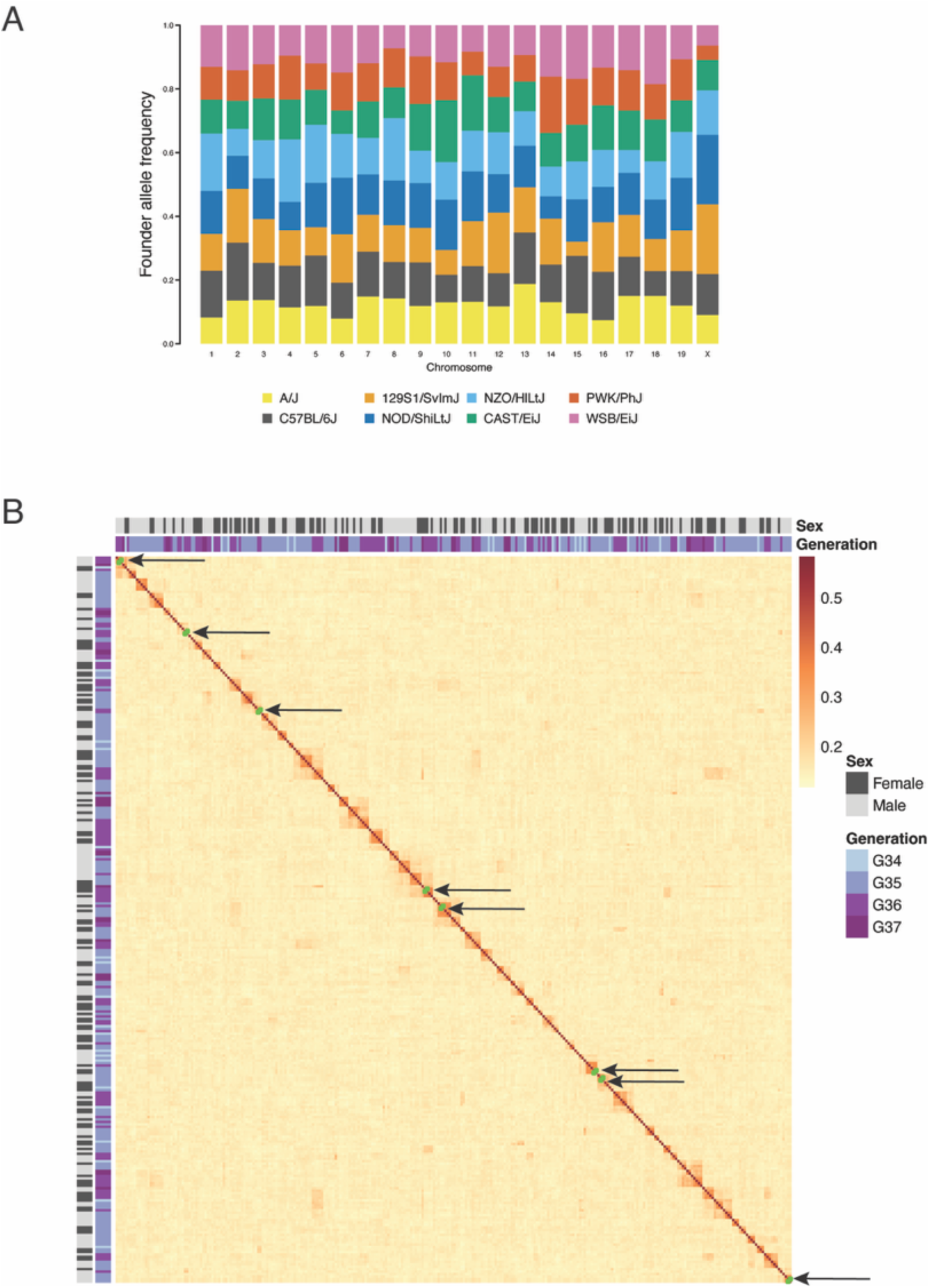
Genetic analysis of the DO iPSC panel. (A) Founder haplotype frequencies across chromosomes were estimated from GigaMUGA genotypes. Contributions from each of the eight DO founder strains are shown for chromosomes 1–19 and X. Deviations from the expected 0.125 founder contribution are summarized in Supplemental Table 2. (B) Pairwise kinship coefficients were calculated from GigaMUGA-derived genotypes to assess relatedness among iPSC lines. Eight pairs of lines showed high kinship (kin_raw > 0.4; highlighted in green and arrows), and one member of each pair was flagged in the final panel (Supplemental Figure 1). All remaining line pairs showed kinship values consistent with non-sibling Diversity Outbred mice.

To investigate relatedness within the DO iPSC panel, we estimated pairwise kinship (K) from genotype probabilities using R/qtl2 [34]. As expected given the random outbreeding scheme of the DO population [6] and our selection of non-sibling animals for derivation of tail tip fibroblasts [19], most pairwise line comparisons showed tightly distributed kinship values equivalent to first cousins (median = 0.151; mean = 0.154), with 95% of pairs below 0.171 and 99% below 0.230 (Supplemental Figure 1A; Fig. 2B). In contrast, 8 pairs of samples exhibited kinship exceeded the expected kinship for siblings (K > 0.50) and approached the value expected for genetically identical samples. These pairs were therefore classified as likely duplicate lines (Fig. 2B, highlighted in green). Five of the 8 pairs were derived from animals with unique IDs, suggesting that duplication occurred during derivation or expansion of the iPSCs (one member of each pair is flagged as a potential duplicate in Supplemental Table 1).

To characterize genetic variation in the panel and assess structure across generations, we calculated the fixation index (FST), inbreeding coefficient (FIS), minor allele frequency (MAF), and heterozygosity. The DO iPSC lines were derived from four successive outbreeding generations (G34–G37), raising the possibility of subtle allele-frequency differences among generations. In a separate multi-generation DO cohort, Tyler et al. (2021) reported FST = 0.022 based on genetically inferred subpopulations [35]. Using generation as the grouping variable, we estimated a comparable value (FST = 0.017), indicating weak differentiation among generations.

To evaluate deviation from Hardy–Weinberg expectations, we calculated the inbreeding coefficient (FIS) from the genotyping data. FIS quantifies the difference between observed and expected heterozygosity, with values near zero indicating concordance with Hardy–Weinberg equilibrium. The panel exhibited a low FIS (0.0097), consistent with minimal deviation from expected heterozygosity. We further examined minor allele frequency (MAF) and heterozygosity and found values consistent with prior reports for DO populations (mean expected:observed heterozygosity = 0.367:0.364; median MAF = 0.278) [36]. Together with the low FST, these metrics indicate that the iPSC panel retains the allele frequency spectrum and heterozygosity of the outbred population from which it was derived.

### Growth and early lineage commitment

We randomly selected 37 cell lines and expanded these over 5 independent rounds to evaluate doubling time (Fig 3A). We calculated the growth-rate constant for each line and compared these by chromosomal sex. XY and XX lines exhibited a mean doubling time of 17.2 and 20.3 hours, respectively, however this sex difference was not statistically significant (p = 0.32, Welch’s Two Sample t-test). These doubling times are similar to those reported for other mouse iPSC lines [37], and to our knowledge, potential sex differences in doubling time have not previously been explored in mouse iPSCs.

**Figure 3.**
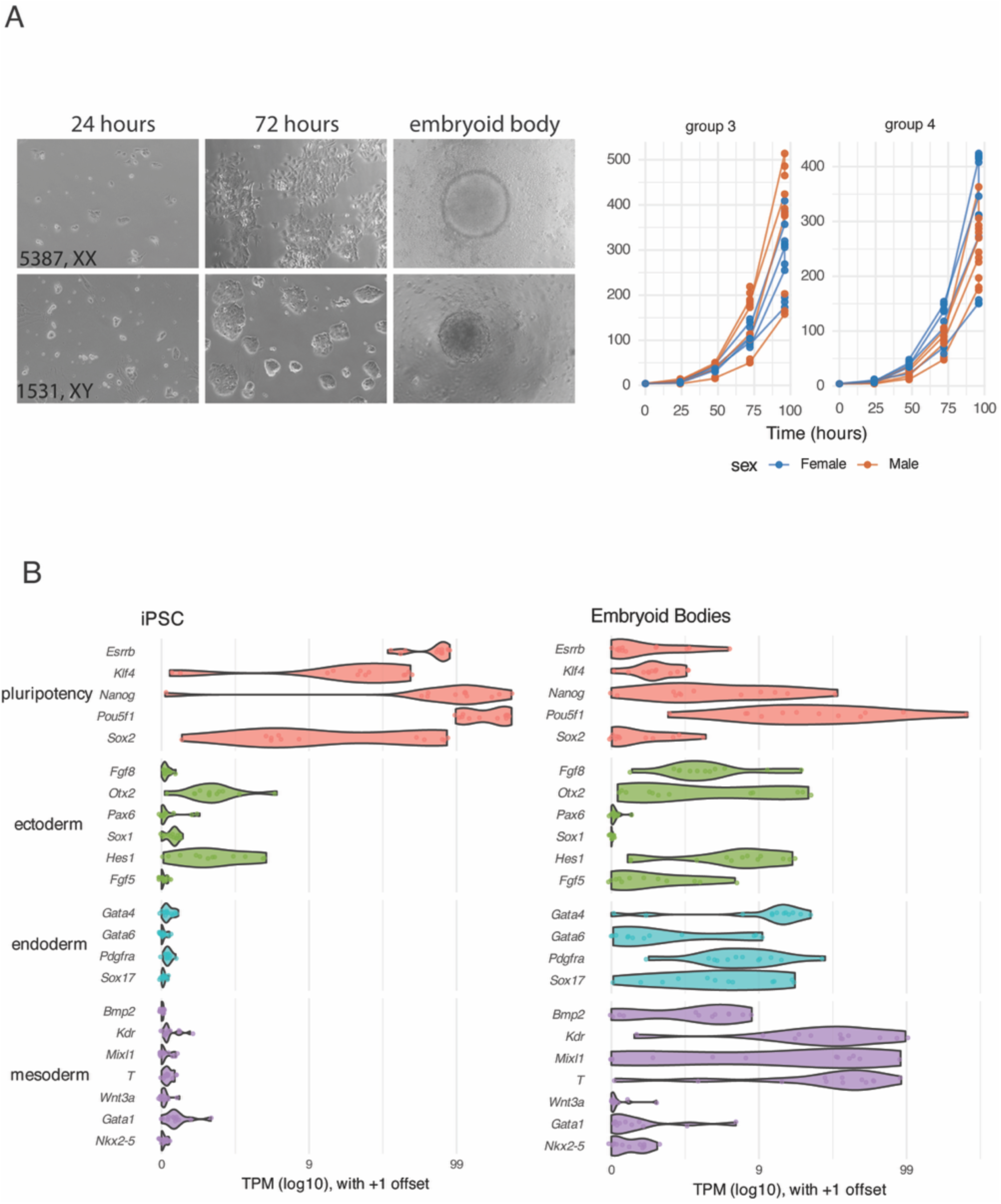
Growth characteristics and embryoid body differentiation of DO iPSC lines. (A) Example of growth profiling of male and female DO iPSC lines. Cells were cultured for 96 hours and cell counts were used to calculate doubling time. Lines were assayed across multiple experimental batches. Representative colony morphologies are shown, including flattened morphology observed in some lines (e.g., line 5387, XX, 72 hours). (B) Embryoid body (EB) formation from 12 randomly selected iPSC lines. All lines formed EBs, with variation in EB morphology, some smooth and spherical and some rough and irregular, depending on the line. (C) Gene expression analysis before and after EB differentiation. RNA was collected from undifferentiated iPSCs and from EBs. Expression of pluripotency markers and early lineage markers representing ectoderm, mesoderm, and endoderm was quantified. Undifferentiated lines showed high pluripotency marker expression and low lineage marker expression, whereas EBs showed reduced pluripotency marker expression and increased lineage marker expression.

To functionally evaluate the DO iPSC lines, we randomly selected 12 lines, balanced by XX and XY for differentiation into embryoid bodies (EBs). RNA was collected before and after differentiation and expression patterns of canonical pluripotency-associated genes and early lineage markers were used to evaluate inter-line variation in molecular signatures of pluripotency and early lineage commitment. Each of the 12 selected lines exhibited high expression of pluripotency-associated genes prior to differentiation. We also detected expression of the epiblast marker *Otx2* in the undifferentiated iPSC lines. Expression of this marker gene is typically observed in primed pluripotent stem cells like mouse EpiSCs and human ESCs / iPSCs, however other markers typically expressed in primed mouse ESCs or EpiSCs were not highly expressed (i.e. *Fgf5*) [38]. In EBs, we detected expression of markers associated with all three early lineages, but with a high variability in their expression across the 12 lines, consistent with previous studies of early differentiation in genetically heterogeneous pluripotent mouse cell lines [15, 17, 18, 20, 21] (Fig 3B).

### Transcriptomic profiles

To explore the transcriptional heterogeneity more comprehensively, we randomly selected 218 cell lines (balanced for sex) for transcriptomic profiling. We quantified the expression of 14,496 genes that were detected in at least 50% of the cell lines. We found high expression of canonical pluripotency-associated genes (*Pou5f1 [Oct3/4], Nanog, Sox2, Esrrb, Klf4*) and as expected, there was inter-line variation in expression of these genes across individual cell lines (Fig 4A). Ectoderm marker *Otx2* showed higher expression than other early lineage markers (Fig 4A), closely matching prior DO mESC datasets [18, 38]. No comparable elevation of mesoderm or endoderm markers was detected, indicating that inter-line variability is confined to the expected pluripotent transcriptional profile without evidence of unexpected lineage specification.

**Figure 4.**
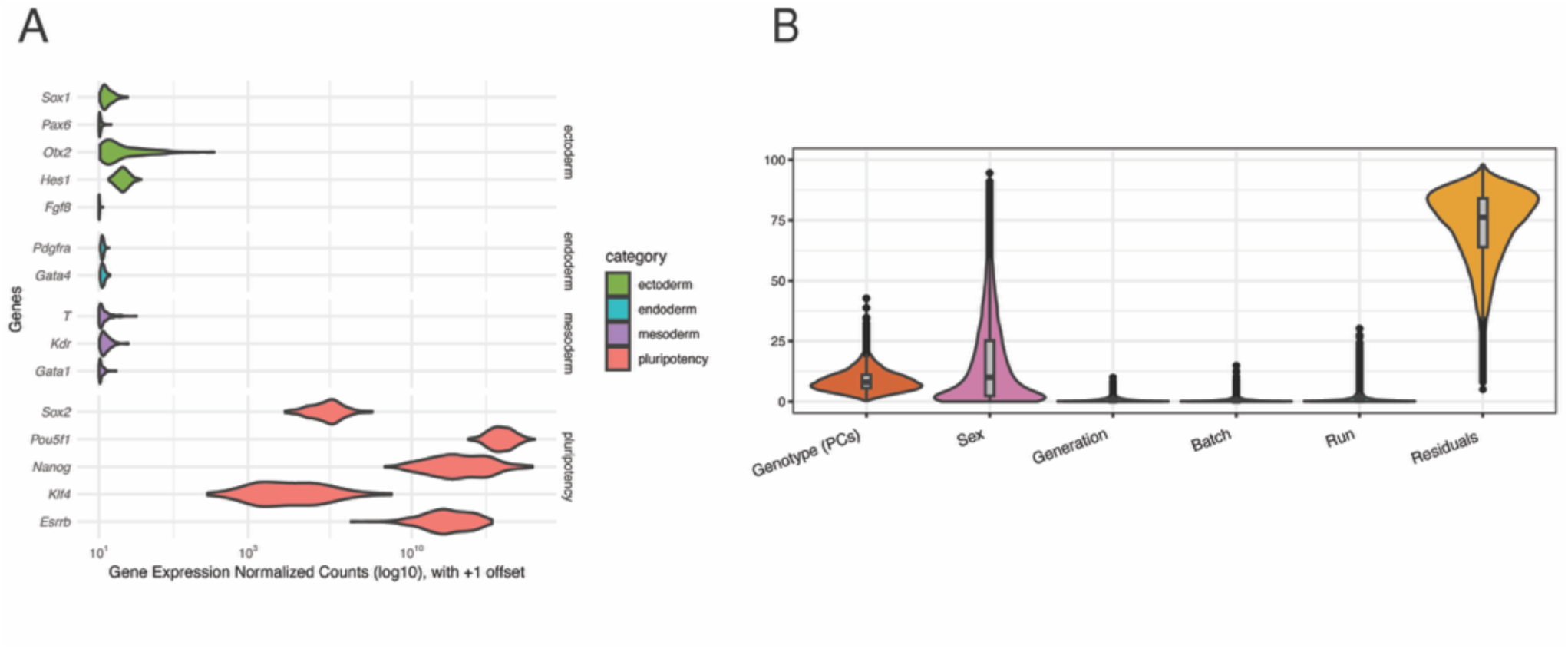
Components contributing to variance in pluripotency gene expression in DO iPSCs. (A) Gene expression of core pluripotency genes and early lineage markers in 218 DO iPSC lines. (B) Variance component analysis showing contributing factors to the variance in gene expression. Genotype is represented by the top 10 principal components informed by genotype probabilities.

### Aneuploidy

While karyotypic analysis of G-banded chromosome spreads remains the gold standard for assessing the degree and type of aneuploidy in cell lines, the size of our iPSC panel and budgetary constraints prevented us from using this approach for all but a few (8) of the total 288 lines. As an alternative, we used our large transcriptomic dataset to infer chromosome-scale aneuploidies in our 218 lines. Chromosome copy number has been previously shown to impact transcript abundance where, for example, a gain in copy number leads to a 1.5x increase on average in gene expression for many of the impacted genes in tissues (e.g. [39]) and in cell lines [28, 40]. This has formed the basis for gene expression-based diagnostics (RT-PCR, RNA-seq) in IVF laboratories and for characterization of cell lines commonly used in biomedical research. We included all autosomal genes expressed in at least 20% of the 218 lines and grouped genes by their encoded chromosome. For each line, we calculated the median expression levels for chromosomes 1-19, and then analyzed the population distribution of these chromosome-level scores to identify outlier lines that were likely trisomic or monosomic [18, 40, 41]. We flagged 33 of 218 iPSCs (15.1%) as having a putative trisomy (28 lines) or monosomy (3 lines), or both (2 lines). As expected, many of the chromosomal aneuploidies were trisomy for chromosome 8 (10 lines) or chromosome 11 (10 lines), both of which have been shown to confer growth advantages in cultured mouse PSCs [42, 43](Supplemental Figure 2).

We compared our gene expression-based aneuploidy calls to karyotyping results for two lines that were profiled by both methods. Expression-based profiling correctly identified one of the lines as euploid (sample 1571, XY, karyotyped as 95% euploid cells) and the other line as being trisomic for chromosome 11 (sample 209, XY, karyotyped as 0% euploid cells, 40% + chr 11, 20% + chr 8). Expression-based profiling using our bulk RNA-seq dataset was unable to detect the additional trisomy chromosome 8 that was found in 20% of the cells in sample 209. To our knowledge, a comprehensive comparison of gene expression-based karyotyping to standard cytogenetic karyotyping has not been published so it is difficult evaluate the threshold at which a particular gain or loss of a chromosome might become detectable in cell culture using transcript abundance but based on our analysis it is at least higher than 20%.

### eQTL maps

Several factors influence gene expression variation in genetically diverse cell panels, including sex and genetic background [18, 44, 45]. Before attempting to genetically map expression quantitative trait loci (eQTLs), we used variance component analysis to determine the contribution of genotype (or genetic background) to transcriptional heterogeneity. While residuals (unknown factors) were high, we found that genotype contributed to 9% of the overall variation in gene expression (Fig 4B). To identify the genetic variants that influence transcriptional heterogeneity in the DO iPSC panel, we performed eQTL mapping. We mapped 10,028 significant eQTLs (LOD > 7.78) that influence the expression of 8,675 genes (59% of all expressed genes). Most significant eQTLs were in proximity (within ± 2Mb) to their target gene and are classified as local eQTL (68%); these loci are visible in the strong diagonal band of the eQTL map (Fig 5A). The remaining 32% of significant eQTL map far from their target gene and are classified as distant eQTL; these loci are evident as points off the diagonal in the eQTL map (Fig 5A). The expression of many genes may be influenced by the same distant eQTL and appear as a vertical band in the eQTL map (Fig 5A). We identified seven such eQTL “hotspots” affecting the expression of many genes in our iPSC panel (Fig 5B), four of which are shared with DO ESCs (Supplemental Table 4) [18].

**Figure 5.**
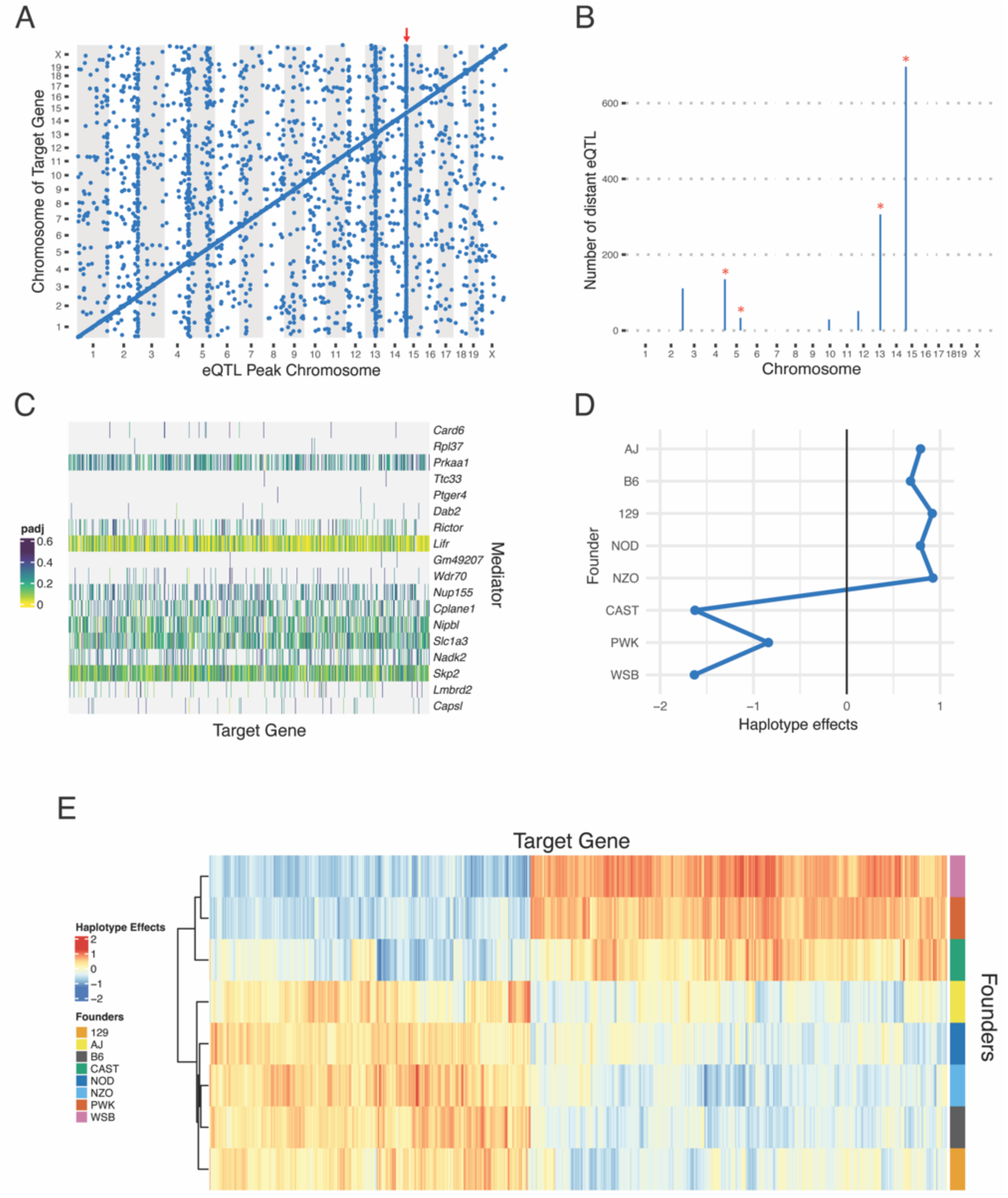
Loci contributing to transcriptional heterogeneity in DO iPSCs. (A) Genomic locations of DO iPSC eQTLs (LOD > 7.78). Red arrow denotes chromosome 15 eQTL hotspot with over 600 distant target genes. (B) Distant eQTL hotspots, where distant is defined as eQTL greater than 2Mbp from its target gene. Asterisks denote hotspots shared with published DO ESC transcriptomics. (C) Heat map of mediation analysis identifying *Lifr* as the strongest mediator for the chromosome 15 hotspot. Potential mediators were identified using the change in LOD score and the associated adjusted pvalues for each mediator-target interaction are represented by color. (D) Haplotype effect at the local eQTL for *Lifr* showing a 3:5 split, with more recently wild-derived strains grouping separately from the classic inbred strains. (E) Heatmap of the haplotype effects at the 696 distant eQTL within the chromosome 15 hotspot showing a 3:5 split between the wild-derived and classic inbred strains.

The largest eQTL hotspot maps to proximal chromosome 15 and influences the expression of 696 target genes (Fig 5B). Consistent with prior eQTL mapping studies in DO mESCs [18], *Lifr* is the most likely mediator gene underlying this shared distal eQTL (Fig 5C). *Lifr* encodes a component of the LIF receptor complex [46] and exhibits a strong local eQTL in the iPSC panel. In contrast to the previously reported 4:4 founder allele pattern in DO mESCs, the *Lifr* locus in iPSCs displays a 5:3 pattern, with the NOD/ShiLtJ allele grouping with the classical inbred founder strains (Fig 5D). This shift is recapitulated across the hotspot target genes (Fig 5E), indicating a coordinated change in allele effects relative to the mESC dataset, where NOD/ShiLtJ clustered with the wild-derived strains [18]. This difference is not attributable to founder representation in the cell population, as allele frequencies at the *Lifr* locus were comparable across lines. Instead, the altered allele effects align with differences in culture conditions between studies. DO iPSCs were maintained in 2i/LIF, which stabilizes ground-state pluripotency and modulates NOD/ShiLtJ-associated expression programs [47], whereas the conditions provide a plausible mechanism for the observed shift in allele effects at the *Lifr* locus and its downstream targets. The chromosome 15 hotspot in iPSCs includes nearly twice as many target genes as reported in mESCs, consistent with increased sample size and sequencing depth improving power to detect trans-eQTL. Together, these results indicate that while the *Lifr*-mediated regulatory axis is preserved across pluripotent systems, its allele-specific effects are sensitive to cellular state and culture conditions.

## CONCLUSIONS

We generated and characterized a panel of iPSCs derived from Diversity Outbred mice and created a biobanked community resource for genetic and functional studies. Using high-density SNP-genotyping, we found that the panel preserves the genetic architecture and heterozygosity of the source population. Functional assays demonstrated pluripotency across lines, with high expression of canonical pluripotency-associated genes and the ability to form embryoid bodies that express markers of ectoderm, mesoderm, and endoderm. Transcriptomic profiling revealed expression heterogeneity across lines, and expression quantitative trait locus mapping identified over 10,000 eQTL, including multiple regulatory hotspots shared with DO embryonic stem cells. These results show that naturally occurring genetic variation shapes transcriptional regulation in mouse iPSCs and establish this DO iPSC collection as a genetically diverse platform for investigating the genetic basis of cellular phenotypes and regulatory variation.

## MATERIALS AND METHODS

### iPSC reprogramming

Diversity Outbred (DO, IMSR_JAX:009376) tail tip fibroblast lines (P3) derived for a previous study [19] were used as starting material for iPSC reprogramming. Fibroblasts were converted to induced pluripotent stem cells through doxycycline induced, ectopic expression of the four reprogramming genes (‘factors’), *Oct4*, *Sox2*, *Klf4*, and *cMyc* (OKSM) delivered via lentiviral transduction. Lentiviruses were produced from 293T packaging cells following transfection with a 6 plasmid system that included pHage2-EF1a-rtTA-ires-Puro (backbone), phage2-tetO-mOKSM (backbone), pHDM-Hpgm2 (gag/pol), pHDM-tat1b (tat), pRC-CMVRev1b (rev), and pHDM-VsB-G (vsv-g), allowing for the assembly of a single inducible polycistronic cassette as previously described [48, 49]. Vectors were a gift from Dr. Matthias Stadtfeld.

Approximately 100,000 DO tail tip fibroblasts were seeded on plastic in 35-mm culture plates in mouse fibroblast growth media (RPMI 1640 media [Gibco, 11875093] containing 1X penicillin/streptomycin [100X, Gibco 15140], 1X GlutaMAX [100X, Gibco, 35050061], 1X non-essential amino acids [100X, Gibco 11140], 0.0005% 2-mercaptoethanol [55 mM, Gibco 21985023], and 10% fetal bovine serum [Gibco, 16141079]) and infected with 15 μl of concentrated virus in the presence of polybrene (5 μg/ml). Medium was replaced after 16 hours with mouse embryonic stem (mES) cell medium (DMEM supplemented with 15% ES cell grade FBS [Gibco, 16141079], 1X GlutaMAX [100X, Gibco, 35050061], 1X penicillin/streptomycin [100X, Gibco, 15140], 1X nonessential amino acids [100X, Gibco, 11140], 1X sodium pyruvate [100X, Gibco, 11360], 0.1 mM β-mercaptoethanol [55 mM, Gibco 21985023], and 1,000 U/ml leukemia inhibitory factor [Chemicon ESG1106 or 1107], supplemented with 2i [3 uM CHIR99021, Stemgent 04-0004, GSK3 inhibition and 1 uM PD0325901, Stemgent 04-0006, MEK/ERK inhibition]) with daily media changes. To initiate reprogramming ascorbic acid was added to stabilize epigenetic marks [50] along with doxycycline (1 ug/ml, Sigma-Aldrich) to induce expression of the reprogramming factors. Doxycycline and ascorbic acid were removed at day 10 postinfection. iPS colonies were collected 20-25 days postinfection and polyclonal cultures were expanded by plating on mitomycin C-treated MEFs in mES cell medium. Cell lines were frozen at P3 in mESCs media (without LIF or 2i) containing 10% DMSO and 20% FBS. As the iPSC lines were derived, we observed that the numbers of days between P1 and P2 ranged from 2 to 8 days, with a mean of 4.25 days. During reprogramming, we endeavored to retain every line, regardless of its growth rate, to avoid selection of lines with fast growth rates that are typically driven by certain karyotypic abnormalities (see Results, *Aneuploidy*).

Each of the lines were thawed and expanded to P5-P6 in mESCs medium +2i/LIF for characterization and banking. DNA was collected from each line and genotyped using the GigaMUGA platform (Neogen, 550) [32]. Cell culture contaminants were tested using standard bacterial / fungal culture and qPCR for mycoplasma as previously described [25].

### Genetic Analyses

#### Haplotype reconstruction

Genotype and phenotype data were analyzed using the *R/qtl2* software package [34], which is designed for quantitative trait locus (QTL) mapping in multiparent populations. All analyses were conducted in R version 4.4.0 using the default R/qtl2 workflows for the Diversity Outbred population. Raw genotype calls were converted to founder allele probabilities using the corresponding genotypes from the 8 DO founder inbred strains as reference. From the founder allele probabilities, most likely founder diplotypes were reconstructed along the genome for each iPSC line.

#### Founder allele frequency

Founder haplotype contributions were estimated from R/qtl2 founder allele probabilities by integrating founder-specific probabilities across pseudomarker positions using genetic-distance weights. On chromosome X, contributions were scaled by sex-specific copy number. Founder contributions were summed across individuals to obtain chromosome-specific and genome-wide founder counts, which were converted to frequencies. Deviation from the DO expectation of 1/8 per founder was assessed using χ² goodness-of-fit tests. (i) Per-chromosome: for each chromosome, the 8-element founder frequency vector was tested against p₀ = 1/8; raw p-values were adjusted across chromosomes (1–19, X) using Benjamini–Hochberg FDR, reporting both raw and adjusted p-values and a significance threshold of FDR 0.05. (ii) Genome-wide: aggregated founder counts were tested against p₀ = 1/8, and χ² statistic (df = 7) and p-value were reported, along with per-founder observed frequency, expected frequency, deviation, Pearson residuals, and χ² components.

#### Kinship

To evaluate kinship, an overall kinship matrix accounting for the complex structure of the DO genomes was computed with the diplotype probabilities using R/qtl2. Five additional pairs of iPSC lines formed highly related pairs based on kin_raw estimates (kin_raw>=0.5). This threshold was guided by the expected kinship for near-identical DO samples under inbreeding (0.5 + 1/16) and by the kin_raw values observed for confirmed same-animal pairs, which were slightly below 0.5625. For high kinship pairs, one sample from each pair was flagged for future reference (Supplemental Table 1).

#### Minor allele frequency (MAF), Heterozygosity (H), and Inbreeding (F_IS_, F_st_)

##### Genotypes and preprocessing

GigaMUGA genotypes (A/H/B) were converted to dosages 0,1,2 representing the count of the B allele; missing or no-call entries were set to NA. After sample QC (duplicates/near-clonal pairs removed), analyses used *n*=288. SNPs were filtered at call rate ≥ 0.90 and minor allele frequency (MAF) ≥ 0.01 across the analyzed set. All metrics below were computed on the post-filter matrix of typed SNPs.

##### Allele frequencies and MAF

For each SNP, the B-allele frequency p was estimated as p = mean(dosage)/2 across all samples. Minor allele frequency was defined as MAF = min(p, 1 − p). The distribution of MAF across all retained SNPs was summarized by reporting the mean, median, and selected quantiles. ABH to dosage conversion and summary statistics were implemented in custom code. Thresholds were call rate >= 0.90 and MAF >= 0.01.

##### Expected and observed heterozygosity

Expected heterozygosity under Hardy–Weinberg equilibrium was computed as Hexp = 2 p (1 − p) for each SNP. Observed heterozygosity was computed as Hobs = mean(dosage == 1) for each SNP.

##### Population structure analysis (F_st_)

To quantify population structure in the DO iPSC panel, the fixation index (F_st_, <u>S</u>ubpopulation within the <u>T</u>otal population) was calculated following the approach similar to that described by Tyler et al. (2021) [35] with some modifications. The GigaMUGA genotype matrix (markers × samples) with calls coded as “A”, “B”, or “H” (heterozygous) was used as input. Samples flagged as technical duplicates were removed prior to analysis. Generations (G34–G37) were treated as subpopulations. For each SNP, heterozygosity (π) was calculated as:

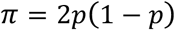

where *p* was the allele frequency across individuals. Overall heterozygosity across all samples (π_T_) and the average heterozygosity across generations (π_S_) were computed. The genome-wide fixation index was then defined as:

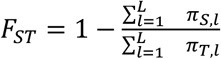

Calculations were performed in R (v4.4) using custom scripts based on the genotype calls and covariate metadata.

##### F_IS_ (Individual within the <u>S</u>ubpopulation, aka inbreeding coefficient)

The within-group (i.e. within-generation) inbreeding coefficient was defined as *F_IS_* = 1 − H_obs_ / H_exp_. The panel-wide value was determined by taking the ratio of the mean observed heterozygosity to the mean expected heterozygosity across SNPs, then computing *F_IS_* from that ratio. FIS per generation was determined by replacing H_obs_ and H_exp_ with group means.

### Aneuploidy

Eight iPSC lines were randomly selected for karyotypic analysis by Cell Line Genetics (https://clgenetics.com/). G-banded karyotypes were used to quantify % euploidy and to identify chromosome-specific aneuploidy. Two of these lines, one predominantly euploid (1571 male, [95% euploid]) and one aneuploid (209 male, [40% + chr 11, 20% + chr 8]) lines were subsequently used for comparison with transcriptome-based karyotyping. Karyotyping by RNA-seq was used to identify chromosome-scale aneuploidies using RNA-seq–derived gene expression profiles (see RNA-seq methods), relative transcriptional dosage across chromosomes were used to infer copy number changes. Raw gene-level count data from 218 undifferentiated iPSC lines were used (32,227 Ensembl-annotated genes × 218 samples). Library sizes were normalized using the Trimmed Mean of M-values (TMM) method as implemented in EdgeR to correct for compositional differences among samples. Genes with counts per million (CPM) > 1 in at least 20% of samples were retained to exclude those with negligible or inconsistent expression. (After filtering: 13859 genes × 218 samples). Normalized counts were transformed to log₂ CPM with a prior count of 1 to stabilize variance and mitigate zero inflation (E_gs_ = log_2_((10×6 x raw counts_g,s_ / library size_s_) +1)). We computed z-scores for each gene using the median and median absolute deviation (MAD) rather than the mean and standard deviation, which are more sensitive to outliers. Each gene’s expression across samples was transformed into a z-score. For each gene *g*, let *x*_%,’_denote the log₂-CPM value in sample *s*. The median expression of a gene across all samples was:

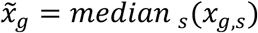

The median absolute deviation (MAD) was defined as:

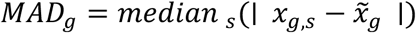

This statistic measures the typical absolute deviation of observations from the median and is less influenced by extreme values than the standard deviation.

The z-score for each gene in each sample was then computed as:

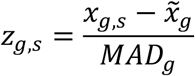

When MAD was extremely small (≤ machine epsilon) or undefined, the median MAD value across all genes was substituted to avoid generating meaningless values.

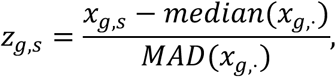

The matrixStats package was used to compute median expression across all samples and MAD.

For each sample, chromosome-level expression scores were calculated as the median of all gene z-scores on that chromosome resulting in one aggregated measure per chromosome per sample. Within each chromosome, distributional parameters were: median chromosome score (*μ*_rob_), MAD (*MAD*_rob_), and standard deviation (*σ*_rob_ = 1.4826 × *MAD*_rob_). Each sample’s chromosome score was evaluated against three empirical thresholds to call aneuploidy: Z-score ≥ 3.5, empirical tail probability ≤ 0.05 (based on the empirical cumulative distribution function, ECDF), absolute deviation (Δ) ≥ 0.25. These thresholds were chosen to be conservative and to minimize false positive calls in the absence of karyotypes consistent with prior expression-based approaches [18, 40]. A chromosome was flagged as putatively aneuploid if all three conditions were met. Quantile–quantile (Q–Q) plots were generated per chromosome using ggplot2 to visualize standardized chromosome scores relative to the normal distribution.

### Growth profiling

iPSC growth profiling was performed according to a previously published method by Udy et al. 1997 [51]. iPSCs were plated in mESM with 2i/LIF at a density of 2 × 10^4^ cells per well on 24-well plates in triplicate (per line), with three wells retained for feeders only on each plate. Triplicate wells were counted over 5 days (0, 24h, 28h, 72h, 96h) using an automated cell counter (Nexcelom). For counting, cells were trypsinized and resuspended in 0.5 ml of mESM. The average cell count of three triplicate wells was used to determine the final count for each time point. 40 DO iPSC lines were randomly selected and divided into 5 experiments (rounds) of 8 lines, 3 lines were discarded due to human error, resulting in 37 lines in the final dataset.

### Embryoid body differentiation

12 lines (6 male and 6 female) were randomly selected for differentiation into embryoid bodies (EB). Cells were resuspended in EB media (mESM without LIF or 2i + bFGF (12ng/mL bFGF added at feed (6 ul/10mL from 20 ng/ul stock) at 750 cells/150ul of EB media and plated onto 96-well ultra-low adherence plates. The plates were centrifuged at 1200rpm for 3 minutes for force aggregation. Aggregated cells were cultured for 48 hours to allow for formation of EBs. EB medium was refreshed at 48 hours by gently replacing 75 ul of media and the EBs were then cultured for an additional 48 hours. EBs were harvested after a total of 4 days in culture (96 hours).

### RNA Sequencing and analysis

#### Embryoid Body Differentiation

Total RNA was isolated using the NucleoMag RNA Kit (Macherey-Nagel) and the KingFisher Flex purification system (ThermoFisher). Undifferentiated iPSCs (iPSC) and embryoid bodies (EB) were homogenized in MR1 buffer (Macherey-Nagel) by vortexing and RNA was isolated according to the manufacturer’s protocol. RNA concentration and quality were assessed using a Nanodrop 8000 spectrophotometer (Thermo Scientific) and the RNA 6000 Pico/RNA Screentape Assay (Agilent Technologies).

Stranded libraries were constructed using the KAPA mRNA HyperPrep Kit (Roche Sequencing and Life Science), according to the manufacturer’s protocol. mRNA was enriched using oligo-dT magnetic beads. Barcoded libraries were created by RNA fragmentation, first and second strand cDNA synthesis, ligation of adapters and barcoding, and PCR amplification. Quality and concentration of the libraries were assessed using the D5000 ScreenTape (Agilent Technologies) and Qubit dsDNA HS Assay (ThermoFisher), respectively, according to the manufacturers’ instructions. Libraries were sequenced 150 bp paired-end on an Illumina NovaSeq X Plus using the 10B Reagent Kit.

Raw RNA-seq reads from induced were processed using the Jackson Laboratory Data Science Nextflow DSL2 Workflows (version 24.10.6, https://github.com/TheJacksonLaboratory/jds-nf-workflows/wiki). Reads were aligned and quantified using EMASE (Expectation Maximization for Allele-Specific Expression) against the *Mus musculus* GRCm39 reference genome [52, 53].

Sequencing quality was assessed using *FastQC* and summarized with the EMASE pipeline’s integrated QC modules. Approximately 45-87 million reads were processed (per sample) with a mean of ∼55.8 million. The mean alignment rate was ∼52% providing ample coverage for gene-level quantification of gene expression.

#### 218 iPSC transcriptomes

Cells were processed, RNA was extracted, and RNAseq reads were generated from 218 cell lines as described above for embryoid body differentiation. Sequencing quality was assessed using FastQC and summarized with the EMASE pipeline’s integrated QC modules. An average of 36 million reads were processed per sample.

Downstream analyses for all RNA-seq datasets were performed in R (version 4.4.0). Gene-level TPM matrices were merged across all samples and annotated with official gene symbols using the biomaRt package. Genes with TPM values below detection across all samples were excluded prior to visualization. TPM values were log-transformed [log₁₀ (TPM + 1)] to stabilize variance and facilitate comparison between conditions. Samples were grouped by cell line and condition (iPSC or EBS), excluding one unpaired EBS sample (line 2524).

Gene expression of the 218 DO iPSC panel subset was normalized per chromosome using upper quartile normalization to account for aneuploidy and then filtered to select genes expressed in greater than 20% of all samples with a median TPM >= 0.5 in the expressed samples. Gene counts were then log-transformed [log_10_ +1 offset] before comparison of known pluripotency and lineage marker comparisons.

Variance component analysis (VCA) was performed to quantify the biological and technical sources that contribute to a proportion of gene expression variability in DO iPSCs. Prior to modeling, tpm-filtered read count values were log-transformed as log_10_ (read counts + 1) to stabilize variance and reduce the influence of extreme values. Since each genotype is only represented once in the dataset, we utilized principal components (PC) identified by eigengenes to simulate genotype. Batch refers to the collection date of the iPSCs while run represents the sequencing date. For each gene, we fit the following mixed-effects model:

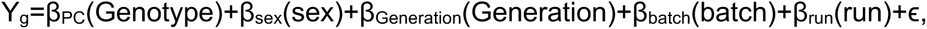

Genes with extremely low overall variance (< 1×10⁻⁵) were excluded prior to modeling, as such genes yield unstable variance estimates. Variance partitioning was carried out using the variancePartition R package [54] and visualized using plotVarPart().

### eQTL Mapping

We performed eQTL mapping on normalized, transformed gene level expression values described above using the QTLretrievR package [55]. Our mapping model included sex as an additive covariate for the eQTL mapping and identified a lod-threshold for genome-wide significance using 1,000 permutations for each of 10 randomly selected genes, establishing a cutoff of LOD > 7.78 for significant eQTL. Distant eQTL were defined as QTL peaks greater than 2Mb from their target gene. Hotspots of distant eQTLs were identified with tallying the significant eQTL (LOD > 7.78) in sliding windows of 50 chromosome-based markers (with 10 marker shift) across the genome andhe top 0.5% of bins with the most eQTL define the hotspots. Mediation of eQTL hotspots was performed using the QTLretrievR package [55] to identify transcripts in that region that are potential causal mediators for the distant eQTL.

### Software

All analyses were performed in R version 4.4.0 (R Foundation for Statistical Computing) on Ubuntu 22.04.4 LTS. Gene expression and related analyses were conducted using Bioconductor packages including limma (3.62.2), edgeR (4.4.2), AnnotationDbi (1.68.0), org.Mm.eg.db (3.20.0), biomaRt (2.62.1), and variancePartition (1.36.3) [54]. Genetic analyses, including genotype probability estimation, kinship calculation, and founder haplotype inference, were performed using the R/qtl2 package [34]. eQTL mapping was performed using the QTLretrievR package (1.2.0.0) [55]. Additional data processing was conducted using dplyr (1.1.4), tidyr (1.3.1), readr (2.1.5), and matrixStats (1.3.0). Visualization was implemented using ggplot2 (3.5.1), ggrepel (0.9.5), and pheatmap (1.0.12). RNA-seq processing and quantification were performed using the Jackson Laboratory Data Science Nextflow DSL2 workflows (https://github.com/TheJacksonLaboratory/jds-nf-workflows, version 24.10.6), with alignment and quantification carried out using EMASE [52, 53]. Figures were assembled using Adobe Illustrator.

## Data Availability

GigaMUGA genotyping data, raw and processed files, and code are available through Figshare https://doi.org/10.6084/m9.figshare.31939140. RNASeq data from the embryoid body differentiation experiments are available from GEO GSE329073. RNASeq data from the eQTL study will be made available through GEO prior to peer review.

## ACKNOWLEDGEMENTS

We gratefully acknowledge the contribution of Genome Technologies and Protein Sciences Services at The Jackson Laboratory for expert assistance and services in support of the work described in this publication. Research reported in this publication was partially supported by the National Cancer Institute under award number *P30CA034196,* a Director’s Innovation Award from The Jackson Laboratory, and the Mouse Mutant Resource Center at The Jackson Laboratory (NIH/ORIP U42OD010921). Vectors were a gift from Dr. Matthias Stadtfeld.

## AUTHOR CONTRIBUTIONS

**Madison Armstrong**, methodology, formal analysis, investigation, writing-original draft, visualization; **Anne Czechanski**, methodology, formal analysis, investigation, data curation, project administration; **Emily Swanzey**, methodology, investigation; **Qionyu Chen**, investigation; **Whitney Martin**, investigation, data curation; **Callan O’Connor**, resources, software; **Catherine Brunton**, investigation; **Selcan Aydin**, methodology, formal analysis, software, supervision; **Hannah Dewey**, formal analysis, software; **Steven Munger**, conceptualization, funding acquisition, formal analysis, supervision, writing – review & editing; **Laura Reinholdt**, conceptualization, funding acquisition, formal analysis, supervision, visualization, writing – original draft, writing – review and editing

## SUPPLEMENTAL FIGURES

**Supplemental Figure 1.**
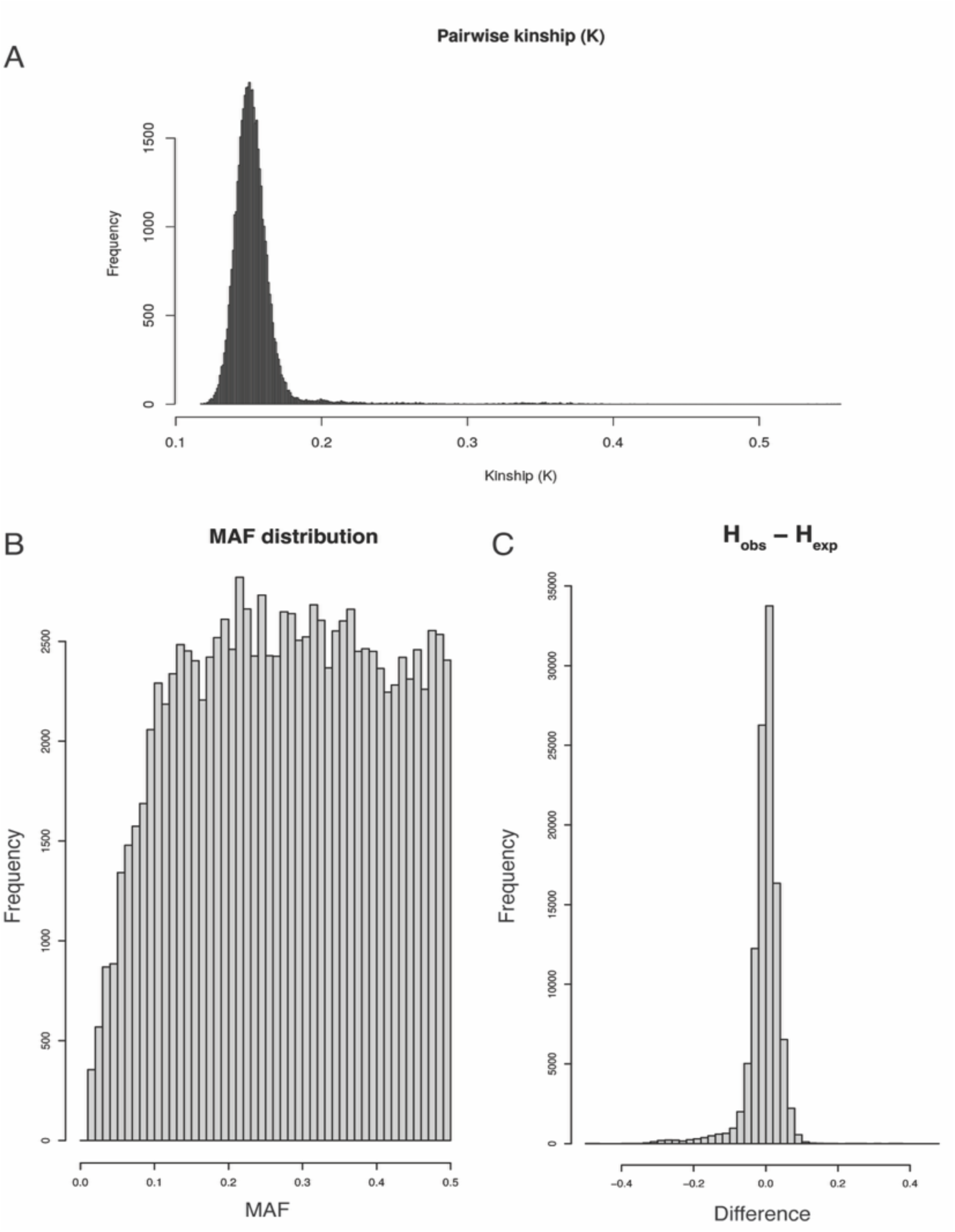
Genetic analysis of the DO iPSC panel. (A) Histogram of kinship values estimated from genotype probabilities across the full DO iPSC panel. Each bar represents the frequency of pairwise comparisons between distinct lines. Most comparisons are tightly centered around K ≈ 0.15 (median = 0.151; mean = 0.154), consistent with the expected relatedness structure of a multiparent Diversity Outbred population. Ninety-five percent of pairs fall below K = 0.171 and 99% below K = 0.230. A small number of pairs are in the extreme upper tail (K > 0.50), approaching the diagonal kinship value expected for genetically identical samples. (B, C) Minor allele frequency (MAF) and heterozygosity (H) were evaluated in the DO iPSC panel and across all measures the population structure of this panel is comparable to a similarly sized genetic mapping cohort of non-sibling DO mice with median MAF = 0.278, mean expected heterozygosity = 0.367 and mean observed = 0.364

**Supplemental Figure 2.**
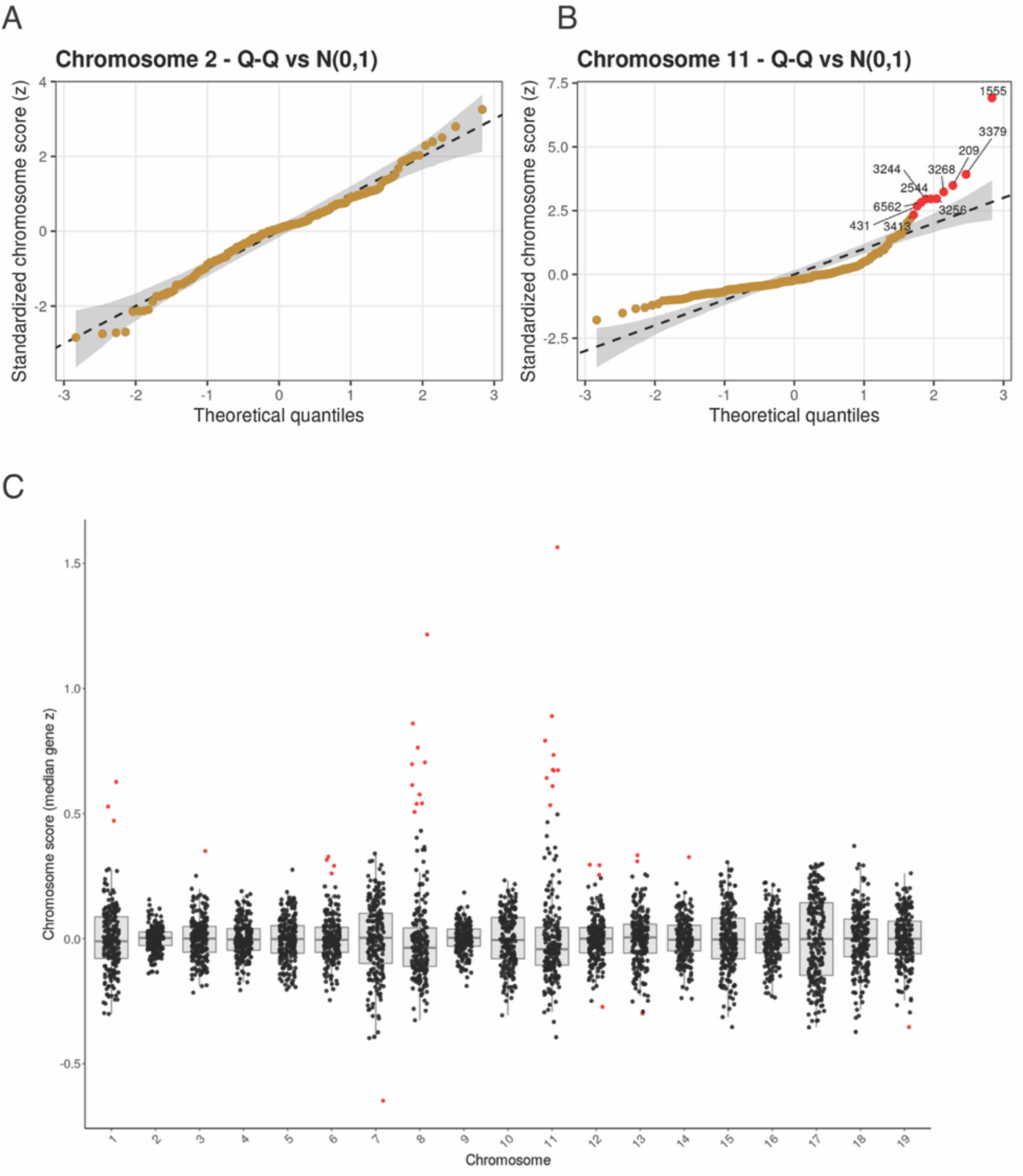
Transcriptome-based aneuploidy analysis. Quantile–quantile (Q–Q) plots of per-sample chromosome expression scores (median gene-level z-scores) were generated for each chromosome using the data from 218 DO iPSC lines. Observed values were statistically compared to the expected normal distribution. Deviation from the diagonal reflects broad chromosome-scale expression shifts. (A, B) Two examples are shown, one for a chromosome without gains or losses (Chromosome 2, A) and one for which there were frequent copy number gains (Chromosome 11, B). (C) To provide an overview for all chromosomes, the chromosome Z score was plotted for each iPSC line. Potential copy number gains were observed for Chromosomes 1, 6, 8, 11, 12, 13, and 14. Potential copy number losses were found for Chromosomes 7, 12, and 19.

## SUPPLEMENTAL TABLES

**Supplemental Table 1. Full table and associated metadata for the DO iPSC resource.** All iPSC lines with associated metadata including unique cell line ID (sample_id), number of days between first and second passage (P1.to.P2.days), passage at first freeze (first.freeze.passage), Jackson Laboratory Biobank ID (BSID), source animal ID (animal.ID), sex of source animal / fibroblast (sex), breeding generation of the source animal (generation), sample number of the tail tip used to generate the fibroblast culture (source.tailtip), source fibroblast ID (fibroblast.name), source cell type (type), passage number of source fibroblast line (fibroblast.passage.reprogram), and flag for cell lines with high kinship scores (duplicate_flag, yes). Fibroblast derivation associated metadata was previously reported in O’Connor et al., 2023 [19].

**Supplemental Table 2. Per-chromosome founder haplotype frequency deviations from expectation.** For each chromosome, deviation of founder haplotype frequency from the expected 0.125 proportion is shown for all eight founders (A/J, C57BL/6J, 129S1/SvImJ, NOD/ShiLtJ, NZO/HlLtJ, CAST/EiJ, PWK/PhJ, WSB/EiJ). Deviations were assessed using a chi-square test against the null hypothesis of equal founder contribution. Raw p-values from Chi-Square (p_raw) and Benjamini–Hochberg adjusted p-values (p_BH) are reported. Columns signif_p and signif_BH indicate nominal and FDR-adjusted significance, respectively. For significant chromosomes, the founder with the largest positive deviation (f1, dev1) and largest negative deviation (f2, dev2) are listed.

**Supplemental Table 3. Chromosome-scale aneuploidies identified by RNA-seq–based karyotyping.** Samples with one or more chromosome-scale copy number alterations inferred from transcriptomic dosage analysis are listed. For each sample_id, the total number of flagged chromosomes (n_chrom_flagged) is reported, along with the specific chromosomes affected. Gains are denoted by g and losses by l.

**Supplemental Table 4. Genomic regions associated with distant eQTL signals.** DO iPSC hotspot information, including genomic locations, number of associated eQTL (LOD > 7.78 or suggLOD > 6.5), and if the hotspot is shared with previously published mESCs transcriptomics

